# Analysis of the risk and protective roles of work-related and individual variables in burnout syndrome in nurses

**DOI:** 10.1101/517383

**Authors:** Mªdel Carmen Pérez-Fuentes, Mªdel Mar Molero Jurado, África Martos Martínez, José Jesús Gázquez Linares

## Abstract

*Burnout* syndrome is a phenomenon that is becoming ever more widespread, especially in workers who have heavy workloads and time pressures, such as nurses. Its progression has been shown to be related to both individual and work-related variables. The objective of this study was to examine the risk and protective roles played by work-related and personal variables, both sociodemographic and psychological, in the development of burnout in nurses. The sample was made up of 1236 nurses. Exploratory tests were performed to understand the relationships between burnout and the other variables, as well as a binary logistic regression to understand their roles in the incidence of this syndrome. Lastly, a regression tree was constructed. The results showed that the sociodemographic variables examined in the study were not related with levels of burnout in nurses. However, certain work-related variables were, such as spending more time with colleagues and patients, and reporting good quality relationships with colleagues, superiors, patients and their families, exhibiting a significant, negative relationship to the presence of burnout. Of the psychological variables, the stress factors conflict-social acceptance, and irritability-tension-fatigue, along with informative communication were found to be risk factors for the appearance of burnout in nurses, in contrast to the communication skills factor, empathy, and energy-joy, which exercised a protective function. The irritability-tension-fatigue factor was the best predictor for the appearance of burnout in nurses.

## Introduction

In recent years we have witnessed increased concern in the healthcare community about workplace performance which has led to numerous studies into its related pathologies and characteristics [1].

*Burnout* syndrome is a phenomenon that is becoming ever more widespread in professionals all over the world, especially in healthcare workers [2], as also occurs in other fields, such as caregiving [3]. It is important because of its consequences, both to the individual and to the workplace [4-5].

### Individual risk and protective factors in the development of burnout syndrome in nurses

Research in this field has established two distinct type of variables related to risk and protective factors associated with the development of *burnout* syndrome. There are personal variables, including sociodemographic and psychological variables, and organizational variables related to nurses’ work environments [6].

Various authors have indicated that sociodemographic characteristics such as sex, age and civil status do not seem to influence the development of burnout syndrome in nurses [7-8]. However, other research indicates that having at least one child, and living with a stable partner reduces the risk of developing burnout [9]. This may be because normally this family circumstance is less common in younger workers who, given their relative inexperience, can suffer higher levels of exhaustion than older workers who have children [10].

Other studies note that living with a partner or having children is a risk factor for developing burnout due to family pressure. Nonetheless, this variable stops being a risk factor in those who work shorter hours [11], as workplace and family expectations no longer produce the same conflict [12].

### Work related factors and burnout syndrome in nursing

Work related characteristics, such as flexible working arrangements, have demonstrated a protective capacity against burnout, as they allow the employee to manage the demands of the job and their own needs [13-14]. In nursing it is common to be pressed for time which gives little opportunity for emotional or physical recovery, and nurses who report lower levels of being pressed for time in their working day exhibit lower levels of burnout [15]. Having limited time can affect important aspects of nursing, such as listening to patients and dealing with their families [16]. Active listening along with empathy, respect and self-knowledge are the requirements for a good therapeutic relationship [17].

Other factors, such as having a fixed contract and longer time in post have been associated with higher levels of burnout in nurses [18]. However, other studies have indicated that, although nurses with more than two decades on the job show higher levels of burnout, it has also been observed in nurses who have been working for eight years or less [19].

### Self-efficacy, self-esteem, communication skills and burnout in nurses

In the organizational context more and more attention is being paid to the study of individual differences and resources as risk and protective factors against stress and work related exhaustion [20] For example, high levels of self-esteem and an appropriate level of self-efficacy are protective factors against burnout [21-22] because what these two variables foster in nurses who have not yet developed burnout may be key in its prevention. Other variables, such as standards related to patient care, are mediating variables in the appearance of burnout in workplaces where there is a heavy workload [23].

Research into burnout has always recognized the central role of social relationships in the development and resolution of this syndrome [2, 24]. Within nursing, and in contrast to other healthcare specialties, there are certain non-technical skills, such as communication, which are central to good job performance [25]. Communication skills in nurses help to protect from and reduce the effects of burnout [26]. The perception of appropriate, effective communication with the patient by healthcare professionals has been related to decreased exhaustion in them and better satisfaction in the patient [27]. In line with that, there is research which indicates that good communication and relationships between team members and their superiors is a protective variable against burnout [28].

In the area of social relationships, the role of emotional intelligence is also notable when responding to and dealing appropriately with overwhelming situations [29-32]. Healthcare professionals’ ability to regulate and manage their feelings is a factor which influences the appearance of burnout [33], especially given the close contact nurses have with patients in intensely emotional situations [34]. An individual’s capacity to manage their emotions, keep calm, and contain empathy and distress allows them to think more clearly, which leads to better patient care [35]. This is why promoting practices which empower nurses to face high-stress situations in an emotionally effective manner is a priority in areas which are especially affectively intense, such as pediatric oncology or palliative care [36-38].

### Stress and burnout in nurses

Workplace stress is a key variable in the development of burnout, as it may appear as a result of a long period of stress on the job. This is why vulnerability to stress is a risk factor for developing burnout [39-40]. Nurses who report facing more stressful situations at work score more highly in burnout than those who report more relaxed activity [41]. In addition, nurses facing hostile and aggressive situations with patients, and feelings of fear and insecurity have been related to higher levels of burnout [42]. Equally, verbal abuse, bullying, threats and physical violence from patients’ families, colleagues, superiors or other specialists have also been linked to diminished job satisfaction and increased burnout and absenteeism [43].

### Objective

Given the above, the objective of this study was to analyze the protective or risk role of job-related variables (such as the type and length of contract, and the quality of interpersonal relationships in the workplace) and personal variables, both sociodemographic and psychological (including social support, communication skills, emotional intelligence, perceived stress, self-esteem and general self-efficacy), on the development of burnout in nurses.

## Method

### Participants

The sample was made up of 1236 nurses aged between 21 and 57, with a mean age of 31.50 (*SD=*6.18). The majority of the sample (84.5%, *n*=1044) were women with a mean age of 31.65 (*SD*=6.23). The remaining 15.5% (*n*=192) were men with a mean age of 30.71 (*SD*=6.17).

Just over half (55%, *n=*680) were single, while 42.1% (*n=*520) were married or in stable relationships. 2.8% (*n=*34) were separated or divorced, and the remaining 0.2% (*n=*2) were widowed. A little over two thirds (68.9%, *n*=852) of the participants had no children, 14.5% (*n*=179) had one child, 13.2% (*n*=163) had two children, and the remaining 3.3% (*n*=41) had three or more.

At the time of the study, 69.3% (*n*=857) worked on short-term or temporary contracts and 30.7% (*n*=379) were employed on permanent contracts.

Almost a third of the nurses (32%, *n*=396) worked on general wards, 21.9% (*n*=271) were emergency room staff, 11.4% (*n*=141) worked in intensive care, 10,7% (*n*=132) in surgical theatres, 2,3% (*n*=28) worked in outpatient settings, 4% (*n*=50) in mental health and the remaining 17,6% (*n*=218) indicated that they worked in other areas.

### Instruments

We used an *ad hoc* questionnaire to collect sociodemographic data. It also included questions about job-related variables such as the type of contract, the amount of time spent each day interacting with colleagues, superiors, patients and families, and the quality of those relationships.

The *Brief Burnout Questionnaire (CBB*) [44] was used to evaluate this syndrome in the nurses. The instrument consists of 21 items in three blocks corresponding to the precursors, elements and consequences of burnout. Despite the objective of the questionnaire being the overall evaluation of the process of professional exhaustion, it addresses the factors proposed in the model from Maslach & Jackson [45] and the components that precede burnout and go along with it. The reliability of the instrument in the sample in our study was 0.87.

Self-efficacy was evaluated via the *General Self-efficacy Scale* [46]. This scale evaluates the feeling of personal competency to deal with stressful situations via 10 items with 4-point Likert-type scale responses. Cronbach’s alpha for the instrument in our study was 0.90.

The *Self-esteem Scale* [47] evaluates an individual’s satisfaction with themselves. It has 10 items, with responses from 1 (“strongly agree”) to 4 (“strongly disagree”) in a Likert-type scale. The reliability of the scale in our study was 0.82.

We evaluated social support using the *Perceived Social Support Questionnaire (CASPE*) [48] This is made up of nine items which determine whether the subject has a partner and the quality of the relationship, social contact and interaction, family relationships and participation in social groups. The first seven items are Likert-type scales with four response options, the eighth item has 5 response options and the final item is answered yes or no. Cronbach’s alpha for this instrument was 0.81.

For the evaluation of emotional skills in the sample, we used the *Brief Emotional Intelligence Inventory* (EQ-I-M20) [49]. This tool has 20 items in five subscales with Likert-type responses. Cronbach’s alpha for each subscale in our study were 0.91 in intrapersonal, 0.72 for interpersonal, 0.82 for stress management, 0.91 in adaptability and 0.88 in the general mood subscale.

We used the *Communication Skills Scale for Healthcare Professionals (EHC-PS)* [50]. This instrument has 18 items with 6-point Likert-style responses. The items are grouped into 4 dimensions: Informative Communication (referring to how clinical information is obtained from or given to patients), which had a Cronbach’s alpha of 0.7 in our study; Empathy (the capacity to understand patients’ feelings, actively listening and responding with empathy), with an alpha of 0.9; Respect (evaluating politeness in the relationship with the patient), with reliability of 0.87 in our study; and Social Skills (the ability to be assertive and socially competent in the clinical relationship), with an alpha of 0.52.

We used the *Perceived Stress Questionnaire* (*Cuestionario de Estrés Percibido*: CEP) [51] to evaluate stress, via 30 statements. The subject reads the statements and indicates the extent to which they reflect their own life through a 4-point Likert-type scale. It is divided into six factors: Tension-instability-fatigue, with an alpha of 0.75; Energy-Joy, with an alpha of 0.80; Overburden, with an alpha of 0.74; Conflict-Social Acceptance, with an alpha of 0.66; Fear-Anxiety with an alpha of 0.57; and Self-realization Satisfaction, with an alpha of 0.62.

### Procedure

The study was approved by the University of Almería Bioethics Committee. Participation in the study was voluntary, and the participants were informed of the study aims, and assured of the anonymity and confidentiality of their responses.

The questionnaires were self-administered and included control questions to check for participants answering randomly. The questionnaires were completed online which took 20-25 minutes. At the beginning of each questionnaire, respondents were given information about how to answer the questionnaire and the type of response in each test.

### Data analysis

First, we present the data on burnout syndrome looking at sociodemographic and work-related variables through frequency analysis. In order to explore the relationship between the variables, we carried out a correlational analysis for the continuous quantitative variables, and the Student *t* test and ANOVA for the categorical variables.

Following that, we performed a binary logistic regression using the introduction method. To that end, the dependent variable (burnout) was dichotomized, the cutoff point was chosen as 25 points based on our assessment of the diagnosis of burnout syndrome. A person scoring over 25 points would be considered to be suffering from the syndrome [44]. The following predictor variables were used: general self-efficacy, overall self-esteem, emotional intelligence (intrapersonal, interpersonal, stress management, adaptability, mood), communication skills (empathy, informative communication, respect, social skills), perceived stress (conflict social acceptance, overburden, irritability-tension-fatigue, energy-joy, fear-anxiety, self-realization-satisfaction), and perceived social support. Lastly, we constructed a regression and classification tree using the CHAID (Chi-Square Automatic Interaction Detector) method.

Treatment and analysis of data was done using the SPSS statistical package version.23 for Windows.

## Results

### Burnout, sociodemographic variables and workplace characteristics

Through the frequency analysis, looking at the presence or absence of burnout syndrome we see that 17.7% (*n*=219) of the nurses scored 25 or higher, as opposed to the 82.3% (*n*=1017) who scored lower. Of those who were affected by burnout, 19.6% (*n*=43) were men and 80.4% (n=176) were women. When we examined the dependent burnout variable without dichotomizing it, we did not find statistically significant differences in the mean scores (*t*=1.03; *p*=.30) between men (*M*= 20.56; *SD*= 5.27) and women (*M*= 20.17; *SD*= 4.75). No differences were seen in burnout related to civil status (*F*=.36; *p*=.77).

We performed correlational analysis in order to examine the relationship between burnout scores and the continuous quantitative variables. We did not find any significant relationship between burnout and age (*r*=.01; *p*=.57) or number of children (*r*=.00; *p*=.99).

With regard to work-related variables, such as the percentage of the workday spent with colleagues, superiors, patients or patients’ families, we found negative correlations between burnout score and the amount of the work-day spent with colleagues (*r*=-.05; *p*<.05) and patients (*r*=-.08; *p*<.01). We found negative correlations between burnout and all cases of evaluation of the quality of relationships in the workplace: the quality of relationships with colleagues (*r*=-.19; *p*<.001), superiors (*r*=-.22; *p*<.001), patients (*r*=-.20; *p*<.001), and patients’ families (*r*=-.23; *p*<.001).

Another work-related variable is the pattern of shifts worked (rotation, 24 hours, nights, mornings/evenings). On applying the ANOVA test, we found no statistically significant difference between the groups (*F*= 2.00; *p*=.11). However, we did find differences according to the type of contract. Nurses with permanent contracts (*M*=21.26; *SD*=5.04) had a higher mean score in burnout (*t*= −5.00; *p*<.001) than those on discontinuous contracts (*M*=19.78; *SD*=4.68).

### Psychological variables and burnout

Table 1 shows that burnout is negatively correlated with all of the factors of emotional intelligence (Intrapersonal: *r*= -.10; *p*<.001; Interpersonal: *r*= -.15; *p*<.001; Stress management: *r*= -.26; *p*<.001; Adaptability: *r*= -.16; *p*<.001; Mood: *r*= -.26; *p*<.001). With respect to the components of perceived stress, burnout positively correlated with conflict-social acceptance (*r*=.46; *p*<.001), overburden (*r*=.36; *p*<.001), irritability-tension-fatigue (*r*=.49; *p*<.001), fear-anxiety (*r*=.36; *p*<.001), and self-realization-satisfaction (*r*=.18; *p*<.001), and negatively correlated with energy-joy (*r*= -.47; *p*<.001).

**Table 1.** Correlations between burnout and variables of emotional intelligence, perceived stress, communication skills, self-efficacy, self-esteem and perceived social support

|  |  | Burnout Syndrome |
| --- | --- | --- |
| Emotional Intelligence | Intrapersonal | -.10*** |
|  | Interpersonal | -.15*** |
|  | Stress management | -.26*** |
|  | Adaptability | -.16*** |
|  | General mood | -.26*** |
| Perceived stress | Conflict-social acceptance | .46*** |
|  | Overburden | .36*** |
|  | Irritability-tension-fatigue | .49*** |
|  | Energy-Joy | -.47*** |
|  | Fear-Anxiety | .36*** |
|  | Self-realization - satisfaction | .18*** |
| Communication Skills | Empathy | -.28*** |
|  | Informative communication | -.15*** |
|  | Respect | -.23*** |
|  | Social skills | .01 |
| General self-efficacy |  | -.19*** |
| Overall self-esteem |  | -.34*** |
| Perceived social support |  | -.27*** |

\*\*\* the correlation is significant at the level .001.

With respect to communication skills, burnout correlated negatively with empathy (*r*= -.28; *p*<.001), informative communication (*r*= -.15; *p*<.001) and respect (*r*= -.23; *p*<.001).

Table 2 gives the means for each of the dimensions of emotional intelligence, comparing those affected by burnout with those unaffected. Nurses unaffected by burnout scored significantly higher in Interpersonal (t=3.66; p<.001; *d*=.27), Stress Management (*t*=6.56; *p*<.001; *d*=.49), Adaptability (t=3.77; p<.001; *d*=.28), and Mood (t=7.57; p<.001; *d*=.56) than those affected by it.

**Table 2.** Emotional intelligence, perceived stress, communication skills, self-efficacy, self-esteem and perceived social support. Descriptive statistics and *t* test according to presence of burnout syndrome

|  | Burnout syndrome |  |  |  |  |  | *t* | *p* |
| --- | --- | --- | --- | --- | --- | --- | --- | --- |
|  | No |  |  | Yes |  |  |  |  |
|  | *N* | *Mean* | *SD* | *N* | *Mean* | *SD* |  |  |
| Intrapersonal | 1017 | 10.66 | 1.93 | 219 | 10.29 | 2.65 | 1.85 | .065 |
| Interpersonal | 1017 | 12.40 | 1.85 | 219 | 11.88 | 2.08 | 3.66*** | .000 |
| Stress management | 1017 | 13.36 | 2.23 | 219 | 12.25 | 2.40 | 6.56*** | .000 |
| Adaptability | 1017 | 11.77 | 2.05 | 219 | 11.19 | 2.13 | 3.77*** | .000 |
| General mood | 1017 | 12.73 | 2.22 | 219 | 11.47 | 2.32 | 7.57*** | .000 |
| Conflict-social acceptance | 1017 | 11.78 | 2.49 | 219 | 14.73 | 3.40 | -12.10*** | .000 |
| Overburden | 1017 | 9.37 | 2.35 | 219 | 10.77 | 2.40 | -7.94*** | .000 |
| Irritability-tension-fatigue | 1017 | 17.04 | 3.57 | 219 | 21.14 | 4.25 | -13.28*** | .000 |
| Energy-Joy | 1017 | 15.05 | 2.82 | 219 | 12.28 | 2.87 | 13.10*** | .000 |
| Fear-Anxiety | 1017 | 3.66 | 1.30 | 219 | 4.51 | 1.47 | -7.82*** | .000 |
| Self-realization - satisfaction | 1017 | 6.88 | 1.19 | 219 | 7.32 | 1.29 | -4.62*** | .000 |
| Empathy | 1017 | 26.51 | 3.33 | 219 | 24.29 | 4.29 | 7.21*** | .000 |
| Informative communication | 1017 | 28.88 | 3.39 | 219 | 27.63 | 4.30 | 4.03*** | .000 |
| Respect | 1017 | 16.45 | 1.98 | 219 | 15.31 | 2.55 | 6.23*** | .000 |
| Social skills | 1017 | 16.84 | 3.01 | 219 | 16.71 | 3.21 | .54 | .587 |
| General self-efficacy | 1017 | 32.35 | 4.21 | 219 | 30.84 | 4.46 | 4.76*** | .000 |
| Overall self-esteem | 1017 | 33.39 | 4.22 | 219 | 30.48 | 4.34 | 9.19*** | .000 |
| Perceived social support | 1017 | 24.72 | 2.97 | 219 | 22.77 | 3.58 | 7.52*** | .000 |

\*\*\**p*<.001

In terms of perceived stress, those affected by burnout syndrome scored significantly higher in conflict-social acceptance (*t*=-12.10; *p*<.001; *d*=.90), overburden (*t*=-7.94; *p*<.001; *d*=.59), irritability-tension-fatigue (*t*=-13.28; *p*<.001; *d*=.99), fear-anxiety (*t*=-7.82; *p*<.001; *d*=.58), and in self-realization satisfaction (*t*=-4.62; *p*<.001; *d*=.34) than those unaffected by it. In the energy-joy dimension, however, it is those who were not suffering burnout who scored highest, with the difference being statistically significant (*t*=13.10; *p*<.001; *d*=.98).

The results of the analysis of the mean communication skills scores with respect to suffering from burnout or not. In this case there were significant differences between the groups, with those unaffected by burnout scoring more highly in empathy (*t*=7.21; *p*<.001; *d*=.54), informative communication (*t*=4.03; *p*<.001; *d*=.30) and respect (*t*=6.23; *p*<.001; *d*=.46).

Finally, the results of the comparison between the group suffering from burnout and those unaffected by it in self-efficacy, self-esteem and perceived social support. In this case, nurses who were not affected by burnout scored more highly in general self-efficacy (*t*=4.76; *p*<.001; *d*=.35), overall self-esteem (*t*=9.19; *p*<.001; *d*=.69), and perceived social support (*t*=7.52; *p*<.001; *d*=.56).

### Logistic regression model for the presence of burnout: Risk and protective factors

We performed the logistic regression analysis with burnout syndrome as a dependent variable. This had previously been dichotomized producing two categories: those affected by burnout syndrome, 17.7% (*n*=219), and those unaffected by it, 82.3% (*n*=1017).

The following predictor variables were added to the equation: general self-efficacy, overall self-esteem, emotional intelligence (intrapersonal, interpersonal, stress management, adaptability, mood), communication skills (empathy, informative communication, respect, social skills), perceived stress (conflict social acceptance, overburden, irritability-tension-fatigue, energy-joy, fear-anxiety, self-realization-satisfaction), and perceived social support.

Table 3 presents these variables, regression coefficients, standard error of estimation, the Wald statistic with degrees of freedom and associated probability, the partial correlation coefficient and the odds ratio. The odds ratios for each variable indicate that:

a. From the perceived stress dimensions, conflict-social acceptance and irritability-tension-fatigue act as risk factors with respect to the likelihood of suffering burnout. Nurses with higher mean scores in these dimensions would have a higher risk of developing the syndrome. Energy-joy exercises a protective function against burnout.
b. The elements of communication skills which are significantly involved are empathy (a protective factor) and informative communication (a risk factor).

**Table 3.** Results of the logistic regression analysis for the probability of suffering burnout

| Variables | $\beta$ | Std.<br>Err. | Wald | df | Sig. | Exp($\beta$) | 95% CI |
| --- | --- | --- | --- | --- | --- | --- | --- |
| Intrapersonal | .02 | .03 | .38 | 1 | .538 | 1.02 | .95-1.10 |
| Interpersonal | .03 | .06 | .29 | 1 | .586 | 1.03 | .91-1.16 |
| Stress management | .05 | .04 | 1.91 | 1 | .166 | 1.06 | .97-1.15 |
| Adaptability | -.03 | .06 | .26 | 1 | .610 | .96 | .86-1.09 |
| General mood | .02 | .06 | .13 | 1 | .718 | 1.02 | .90-1.15 |
| Conflict-social acceptance | .17 | .04 | 18.74 | 1 | .000 | 1.19 | 1.10-1.29 |
| Overburden | -.01 | .05 | .08 | 1 | .768 | .98 | .88-1.09 |
| Irritability-tension-fatigue | .15 | .03 | 16.25 | 1 | .000 | 1.16 | 1.08-1.26 |
| Energy-Joy | -.19 | .03 | 24.75 | 1 | .000 | .82 | .76-.88 |
| Fear-Anxiety | -.11 | .08 | 1.58 | 1 | .209 | .89 | .75-1.06 |
| Self-realization - satisfaction | -.08 | .08 | .99 | 1 | .319 | .91 | .77-1.08 |
| Empathy | -.19 | .05 | 13.74 | 1 | .000 | .82 | .74-.91 |
| Informative communication | .09 | .04 | 3.95 | 1 | .047 | 1.10 | 1.00-1.21 |
| Respect | -.03 | .07 | .22 | 1 | .632 | .96 | .83-1.12 |
| Social skills | .03 | .03 | .77 | 1 | .379 | 1.03 | .95-1.11 |
| General self-efficacy | .01 | .02 | .36 | 1 | .544 | 1.01 | .96-1.06 |
| Overall self-esteem | -.01 | .02 | .14 | 1 | .707 | .98 | .93-1.04 |
| Perceived social support | -.01 | .03 | .23 | 1 | .628 | .98 | .92-1.04 |
| Constant | -1.74 | 1.30 | 1.78 | 1 | .182 | .17 |  |

In terms of the fit of the model, we see an overall fit (*χ*^*2*^= 295.40; df= 18; *p*<.001) confirmed by the Hosmer-Lemeshow test (*χ*^*2*^= 5.86; df= 8; *p*= .66). Nagelkerke’s *R*^*2*^ indicates that 35% of the variability in the response variable is explained by the logistic regression model. In addition, from the table of classification of cases we estimate the probability that the logistic function is accurate at 85.4%, with a false positive rate of .03 and a false negative rate of .34.

### Multiple linear regression model of burnout, according to employment situation

We found differences in burnout between nurses on permanent contracts and those who worked under short-term, discontinuous contracts. In order to determine the explanatory value of the psychological variables (analyzed in the overall sample above), we constructed models for each of the groups depending on their employment situation via stepwise multiple linear regression analysis, using the employment situation as the selection variable (discontinuous/permanent).

As table 4 shows, the group of nurses in discontinuous employment (69.3%; *n*=857) produced five regression analysis models, with the final model producing the highest percentage of explained variance 35.3% (*R*^*2*^=.35). The validity of the model was determined using the Durbin-Watson *D* statistic, *D*=2.02. The value of *t* is associated with a probability of error of less than .05 in the variables included in the model (energy-joy, conflict-social acceptance, empathy, irritability-tension-fatigue, and social skills). Of those, Energy-joy has the greatest explanatory value. Collinearity is absent between the variables in the model according to the values of the tolerance indicators and VIF.

**Table 4.**
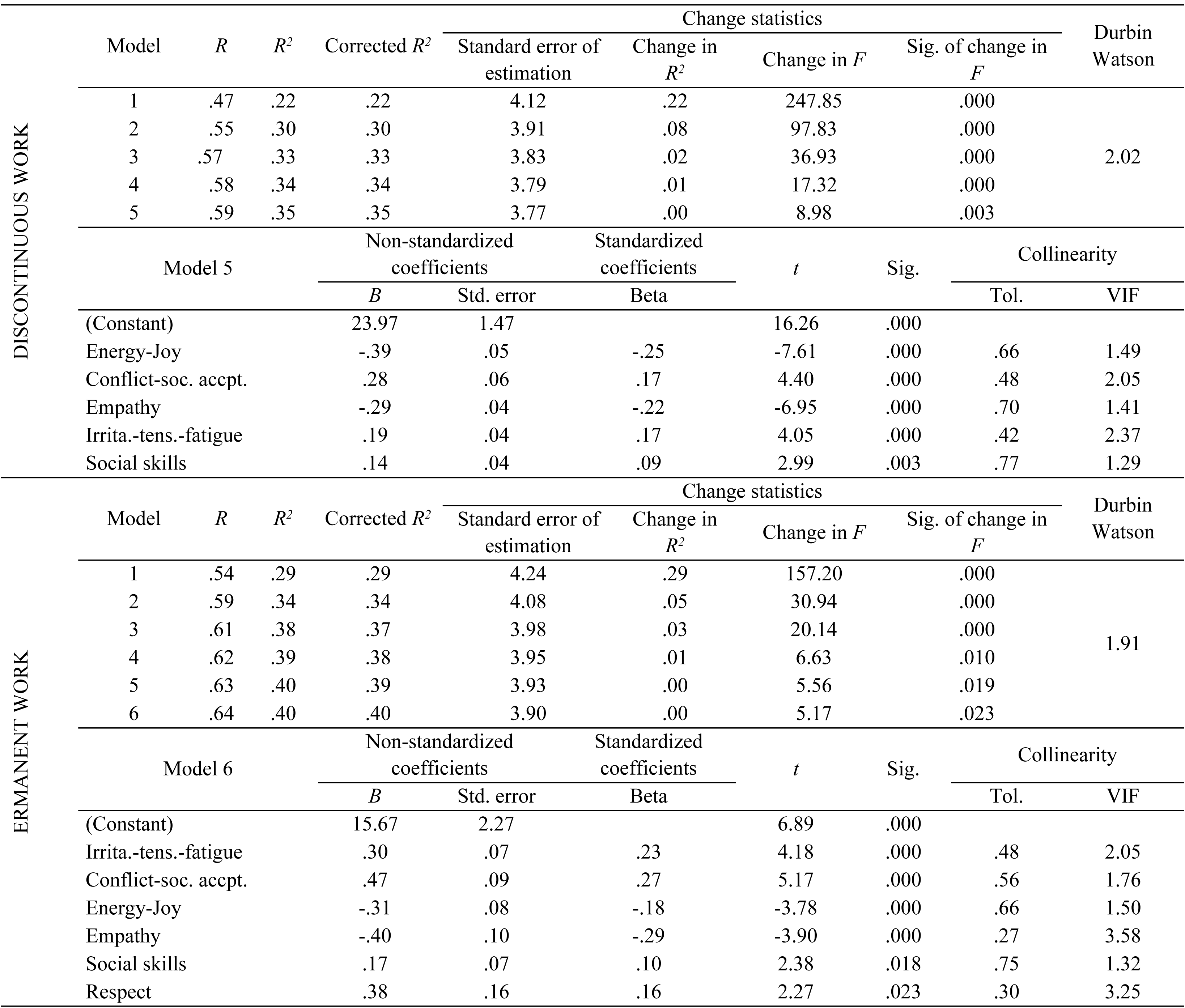
Stepwise multiple linear regression model by employment situation (Discontinuous work: *n*=857; Permanent work: *n*=379)

The regression analyses of the nurses with permanent contracts (30.7%; *n*=379) produced six models. The third model included: Irritability-tension-fatigue, Conflict-social acceptance, Energy-joy, Empathy, Social Skills, and Respect; and explained 40.9% (*R*^*2*^=.40) of the variance. To confirm the model’s validity we analyzed the independence of the residuals using the Durbin-Watson *D* statistic, giving *D=*1.91, which confirmed the absence of positive or negative autocorrelation. The value of *t* is associated with a probability of error of less than in the variables included in the model. The standardized coefficients show that the variable with the greatest explanatory value is conflict-social acceptance. Collinearity is absent between the variables in the model according to the values of the tolerance indicators and VIF.

The decision tree (Figure 1) shows that irritability-tension-fatigue is the best predictor of burnout. Subjects with a high level (>23) of irritability-tension-fatigue exhibited the highest risk of burnout (57.9%). The lowest levels of burnout (96.8%) was found in in subject with very low levels (<14) of irritability-tension-fatigue. Finally the goodness of fit of the model can be seen in its correct classification of 83.7% of the participants.

**Figure. 1.**
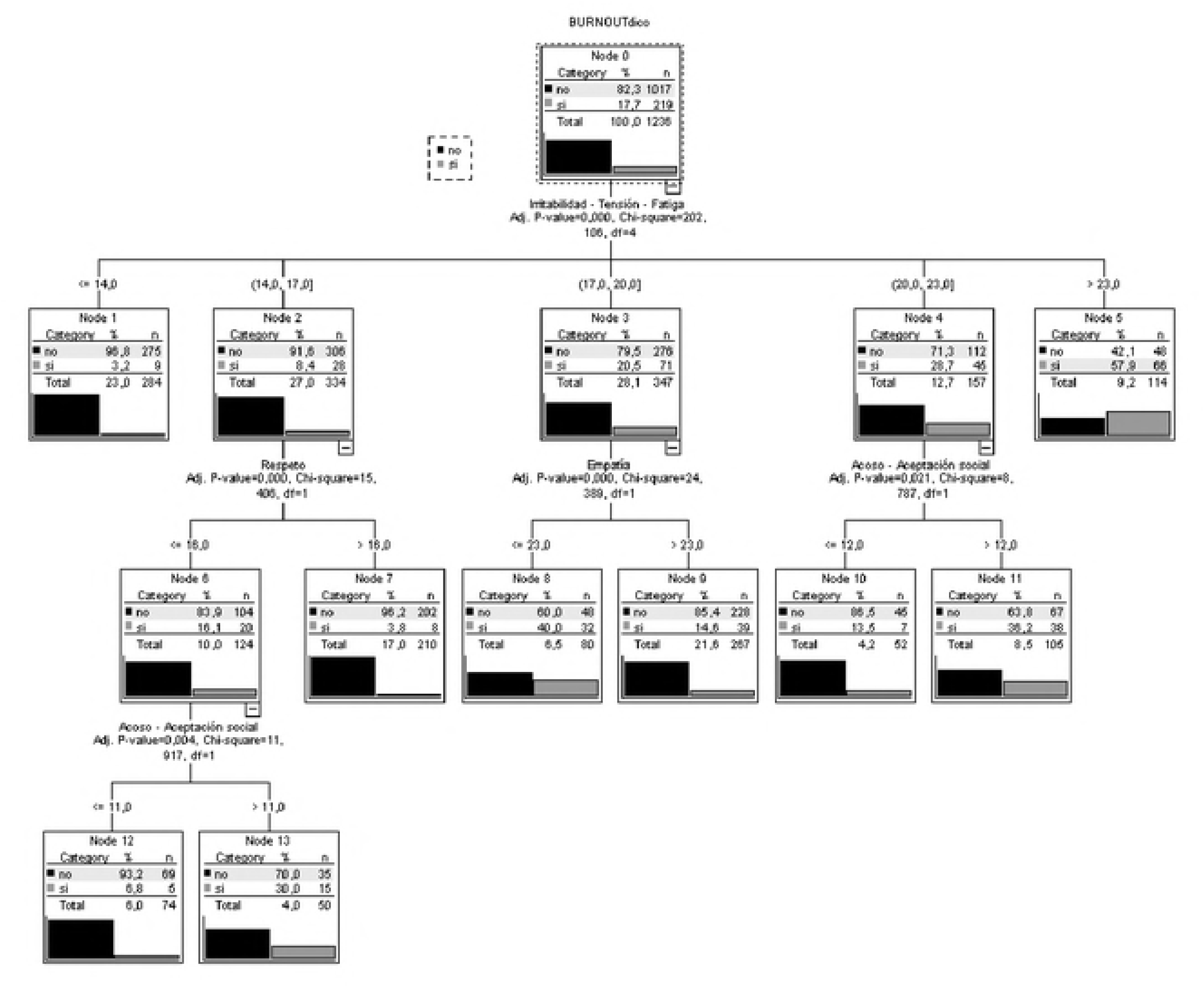
Decision tree according to psychological variables

## Discussion

Our results show that sociodemographic variables such as age, sex, civil status and number of children are not related to levels of burnout in nurses. This is in agreement with other studies in which these individual variables have not exhibited associations with the presence of burnout in these healthcare professionals [7-8].

We did find significant relationships between certain work-related variables and burnout. Spending more time with colleagues and patients, and reporting good quality relationships with colleagues, superiors, patients and their families was found to be negatively related to levels of burnout in nurses. This is in line with the literature, as indicated in research by Lee & Ji [28], having good relationships with members of the team reduces levels of exhaustion. Conversely, negative relationships and conflict are associated with higher levels of burnout [43]. Similarly, the lowest levels of burnout being in nurses who spent more of their work day with colleagues and patients may be due to less time pressure. Those nurses who can spend more time with their patents and colleagues may have less urgency in their daily tasks, which has been associated with lower levels of burnout, as it allows brief periods of physical and emotional recovery [15-16].

In terms of psychological variables, our results show a negative association of burnout with various factors of emotional intelligence [33], self-efficacy, social support [18], communication skills [26] and self-esteem [21]. However, in perceived stress, we found a positive relationship between high levels of burnout and factors associated with conflict, irritability, fatigue, overburden, fear and anxiety [39]. Conversely, energy-joy demonstrated a negative relationship with the presence of burnout, and was highest in those nurses who did not exhibit burnout.

Only some of the psychological variables were found to be protective or risk factors for the development of burnout. Irritability-tension-fatigue in the perceived stress scale, together with informative communication were shown to be risk factors in the appearance of burnout in nurses whereas the empathy factor in communication skills and energy-joy exercised a protective function.

Nurses who work under permanent contracts were seen to suffer higher levels of burnout compared to nurses with discontinuous employment [18]. Once the protective and risk roles of the previous variables had been identified in the overall sample, we carried out a new analysis based on the type of employment. The informative communication variable disappeared for both types of contract, leaving social skills as a risk factor. And in the case of workers with permanent contracts, the respect variable was added to the model as a risk factor.

It may seem strange that informative communication and social skills in nurses, together with respect for patients in nurses on permanent contracts, are risk factors for developing burnout, but it may be due to the following. According to MacPhee et al. [23], exemplary patterns including respect and active communication with empathy that are important in establishing effective therapeutic relationships with patients [17] may also be mediating variables in the appearance of burnout when there is a heavy workload. These variables are not acting as risk factors in and of themselves, but rather are implicated in increased levels of burnout when nurses face high demand with little resources.

## Conclusions

There is a need to do more research evaluating nurses’ workloads, so that we may discover whether communication skills based on respect and assertiveness, and providing optimal information to the patient could be risk factors for exhaustion in these professionals when they face heavy demands.

These findings contribute to the growing body of research for the promotion of improvements in the health of patients and workers, given the negative consequences of burnout to both the workplace and the individuals who suffer from it. Understanding which variables influence its appearance and how they do so will be key elements when it comes to proposing effective prevention and intervention.

## Acknowledgments

The present study was undertaken in collaboration with the Excma. Diputación Provincial de Almería. Part of this work has been developed thanks to the financing of the 2018 Own Research Plan of the University of Almería, for the help for the hiring of research personnel in predoctoral training, granted to África Martos Martínez.

